# The Rodent Electronic Nicotine Delivery System: Apparatus for Voluntary Nose-Only E-Cigarette Aerosol Inhalation

**DOI:** 10.1101/2024.12.21.629932

**Authors:** Amy L. Odum, Mariah E. Willis-Moore, Kiernan T. Callister, Jeremy M. Haynes, Charles C. J. Frye, Lucy N. Scribner, David N. Legaspi, Daniel Santos Da Silva, Aaron L. Olsen, Tadd T. Truscott, Preston T. Alden, Rick A. Bevins, Adam M. Leventhal, Stephen T. Lee, Brenna Gomer, Abby D. Benninghoff

## Abstract

Tobacco use is the leading cause of death globally and in the U.S. After decades of decline, driven by decreases in combusted tobacco use, nicotine product use has increased due to Electronic Nicotine Delivery Systems, also known as e-cigarettes or vapes. Preclinical models of nicotine self-administration can serve as important lodestars in the search for effective intervention and prevention tactics. Current variants of the task have substantial limitations, however. Therefore, we created the Rodent Electronic Nicotine Delivery System, a novel low-cost non-proprietary nose-only preclinical model of nicotine aerosol self-administration. We confirmed that RENDS sequesters nicotine aerosol in the nose port by measuring fine particulate matter (PM < 2.5 microns) generated by e-cigarettes. We also showed that rats robustly self-administer flavored nicotine aerosol, resulting in high blood levels of cotinine (the major nicotine metabolite) and spontaneous somatic withdrawal symptoms. Thus, we provide strong validation of the operation and function of the RENDS, opening the door to an open-source preclinical aerosol model of nicotine self-administration that is relatively low cost. Four existing operant chambers can be retrofitted with the RENDS for less than $325/chamber. All RENDS diagrams and plans for custom designed components are on Open Science Framework (https://osf.io/x2pqf/?view_only=775b55435b8e428f98e6da384ef7889d).

## The Rodent Electronic Nicotine Delivery System: Description and Validation of an Apparatus for Nose-Only E-Cigarette Aerosol Inhalation in Freely Moving Rats

Tobacco use is the leading cause of death globally as well as in the U.S., causing nearly 8 million deaths per year (Reitsma et al., 2021). Nicotine is the primary compound responsible for the extensive use of tobacco (Bevins et al., 2018; Le Foll et al., 2022; United States Department of Health and Human Services [USDHHS], 2014). After decades of decline, nicotine use has increased due to Electronic Nicotine Delivery Systems (ENDS; Cornelius et al., 2022; Cullen et al., 2018). Electronic Nicotine Delivery Systems, also called e-cigarettes or vapes, come in a variety of shapes and sizes, but in general these devices create an aerosol by heating a liquid containing nicotine, most often with flavoring. At the peak, nearly 30% of high school students and over 10% of middle school students in the U.S. used ENDS (Cullen et al., 2019). Current use remains alarmingly high in youth and young adults (Birdsey et al., 2023; Kelsh et al., 2023; Park-Lee et al., 2021), exposing millions of young people to nicotine. Vaping nicotine increases the likelihood of subsequent combusted tobacco use (Center for Disease and Prevention Control [CDC], 2024; Hair et al., 2021; Tashakkori et al., 2023), vulnerability to drug use and misuse (USDHHS, 2016), and harms brain development in adolescents (Castro et al., 2023; England et al., 2015; Leslie, 2020; Yuan et al., 2015). Furthermore, both the constituents of the aerosol as well as nicotine can injure the cardiovascular and respiratory systems (Alanazi et al., 2020; Morris et al., 2021; Park et al., 2019; Qazi et al., 2024; USDHHS, 2016).

To investigate ENDS use, a preclinical model is needed (Corley et al., 2024). Human studies are important but lack control and the ability to isolate key factors contributing to use and misuse (Herman & Tarran, 2020; Jackson et al., 2020). Most prior research on nicotine use in rodents has employed intravenous (IV) self-administration, which allows for strong experimental control, but requires surgery and is limited by relatively short catheter patency duration and the difficulty of implementation with smaller animals (Corley et al., 2024; Pogun et al., 2017). Current research into ENDS use in rodents has employed the whole-body exposure technique in an air-tight “vapor chamber.” In some models, exposure is involuntary (e.g., Frie et al., 2023), whereas in others, rodents can make a response (e.g., a lever press) to introduce aerosol into the chamber (e.g., Smith et al, 2020).

However, the vapor chamber method of pre-clinical ENDS use has substantial limitations. Although rodents can move around freely, and in some preparations voluntarily begin the aerosol presentation, the whole-body exposure method coats the interior of the chamber and the rodent with nicotine-containing aerosol (Chellian et al., 2023). This nicotine then undergoes further absorption through the derma as well as from grooming, confounding measures of drug consumption and physiological impacts (Kogel et al., 2021; Lucci et al., 2020). Furthermore, although a human puff typically lasts a few seconds at most (St. Helen et al., 2018), in a vapor chamber, aerosol presentation is extended many times over the duration of a puff (up to 100 s) while the vacuum system clears the chamber (Lallai et al., 2021; Moussawi et al., 2020). This duration of aerosol inhalation may be stressful, which can change multiple behavioral, physiological, and immunological processes (see Buynitsky & Mostofsky, 2009), and lead rodents to modify their breathing to reduce extended involuntary exposure (Lallai et al., 2021). Furthermore, some studies have failed to show strong control by the operant contingency, finding a low and similar number of responses on the active manipulandum as compared to the inactive manipulandum (Cooper et al., 2024; Marusich & Palmatier, 2023; Smith et al., 2020). This issue may stem from the prolonged and diffuse presentation of the nicotine aerosol, making discrimination of the causal response difficult (Bagdas et al., 2022). Finally, commonly used vapor chambers are also proprietary and cost prohibitive.

We have developed the Rodent Electronic Nicotine Delivery System (RENDS) to address the current limitations of preclinical ENDS models, provide a system with stronger face validity, and reduce barriers to research. The RENDS is a novel, non-invasive, non-proprietary, and relatively low-cost system for snout-only nicotine aerosol self-administration in freely moving rats. The system costs under $325 per chamber to retrofit four existing operant chambers. A custom-designed 3D-printed port records nose pokes and provides access to the aerosol. Here, we describe the RENDS system and present data to validate its function. We completed two main assessments. First, we confirmed that the e-cigarette aerosol cycles exclusively in the nose port, showing that RENDS provides snout-only access. To verify the operation of the nose port, we measured fine particulate matter less than 2.5 microns (PM2.5), found in e-cigarette aerosol and readily measured by air quality monitors (Li et al., 2020; Shearston et al., 2023), with and without the vacuum system of the RENDS in operation. Second, we showed that rats would robustly self-administer flavored e-cigarette aerosol, with concomitant high blood levels of cotinine, the major nicotine metabolite, and accompanying spontaneous somatic withdrawal (Malin et al., 1992). We demonstrated in a reversal design that rats only engaged in robust nose poking when pokes produced flavored nicotine aerosol and related stimuli. These metrics confirm that RENDS is a valid snout-only preclinical model of e-cigarette nicotine aerosol self-administration.

## Validation of the RENDS

### Subjects

We used a convenience sample to reduce animal numbers in the validation phase (NC3Rs, N.D.). Six rats (Charles River Laboratories), four Wistar, and two Long-Evans, were restricted to 85% of the free-feeding growth curve by post-session feeding. Rats lived in same-sex and same-strain pairs in standard Plexiglas rodent enclosures residing in temperature-controlled colony rooms with a 12:12 hour light dark cycle. Experiments were conducted during the light part of the cycle. Colony rooms were in an Association for the Assessment and Accreditation of Laboratory Animal Care (AAALAC) certified facility. Water was freely available in the home cages. The Wistar rats, two male and two female, were approximately 170 days old at the start of the experiment and are hereafter referred to as the nicotine subjects. Nicotine subjects participated in the self-administration, cotinine, and withdrawal assessments. The Long-Evans rats, one male, and one female and hereafter referred to as the control subjects, participated in withdrawal assessments at approximately 120 and 270 days old, respectively. Nicotine rats were 410 days old at the time of withdrawal assessments. All subjects were nicotine-naive but had previous experimental histories with reinforcement schedules of food delivery in operant chambers. All study procedures were approved by the Institutional Animal Care and Use Committee at Utah State University (Protocol #12626).

### Apparatus and Materials

All RENDS system diagrams and plans for custom-made parts are freely available at our Open Science Framework (OSF) link: https://osf.io/x2pqf/?view_only=775b55435b8e428f98e6da384ef7889d. Current costs and parts needed for the RENDS are displayed in Table 1.

**Table 1.** RENDS Components by Description and Cost for Four Chambers.

| Quantity | Component Description | Unit Amount | Total Price (USD) |
| --- | --- | --- | --- |
| 1 | Air compressor |  | 349.00 |
| 4 | Air regulator |  | 67.96 |
| 4 | Ball pump needle | 2 | 3.84 |
| 4 | E-cigarette (Drag X) |  | 179.96 |
| 4 | E-cigarette holder PLA print |  | 13.44 |
| 8 | Photo beam |  | 23.60 |
| 8 | Resin printed port |  | 40.00 |
| 8 | Silicon tubing adapter | 5 | 29.76 |
| 4 | Solenoid |  | 154.00 |
| 8 | Stainless steel adjustable clamp | 10 | 2.48 |
| 1 | Tailonz pneumatic tubing | 50 ft | 18.99 |
| 1 | Vacuum cleaner |  | 109.99 |
| 4 | Vacuum generator |  | 211.20 |
| 1 | Vacuum tubing | 30 ft | 52.87 |
| 1 | Vacuum tubing splitter |  | 4.42 |
|  |  |  | **1,261.51** |

*Note.* This table includes all components and the 2024 costs for four RENDS chambers. One air compressor and one vacuum cleaner can support up to six chambers, which would reduce the per chamber cost.

#### RENDS 4-chamber Set Up

Four identical operant chambers^1^ (Med Associates, St. Albans, VT; ENV-008-B) housed in sound-attenuating cubicles (Med Associates; ENV-022MD) were retrofitted with the RENDS system (see Figure 1). Inside each chamber, two custom 3D printed nose ports (see below for detail) are situated on the left and right side of the front panel 0.5 cm (0.197 in.) above the floor grates. Nose port openings are 3 cm (1.18 in.) above the floor grates. A house light (Med Associates, ENV-215M-LED) is located at the top of the center panel. In each chamber, one nose port is designated as active and delivers nicotine aerosol, and the other is designated inactive and only records nose pokes. Which port is active and which is inactive is counterbalanced across chambers.

**Figure 1.**
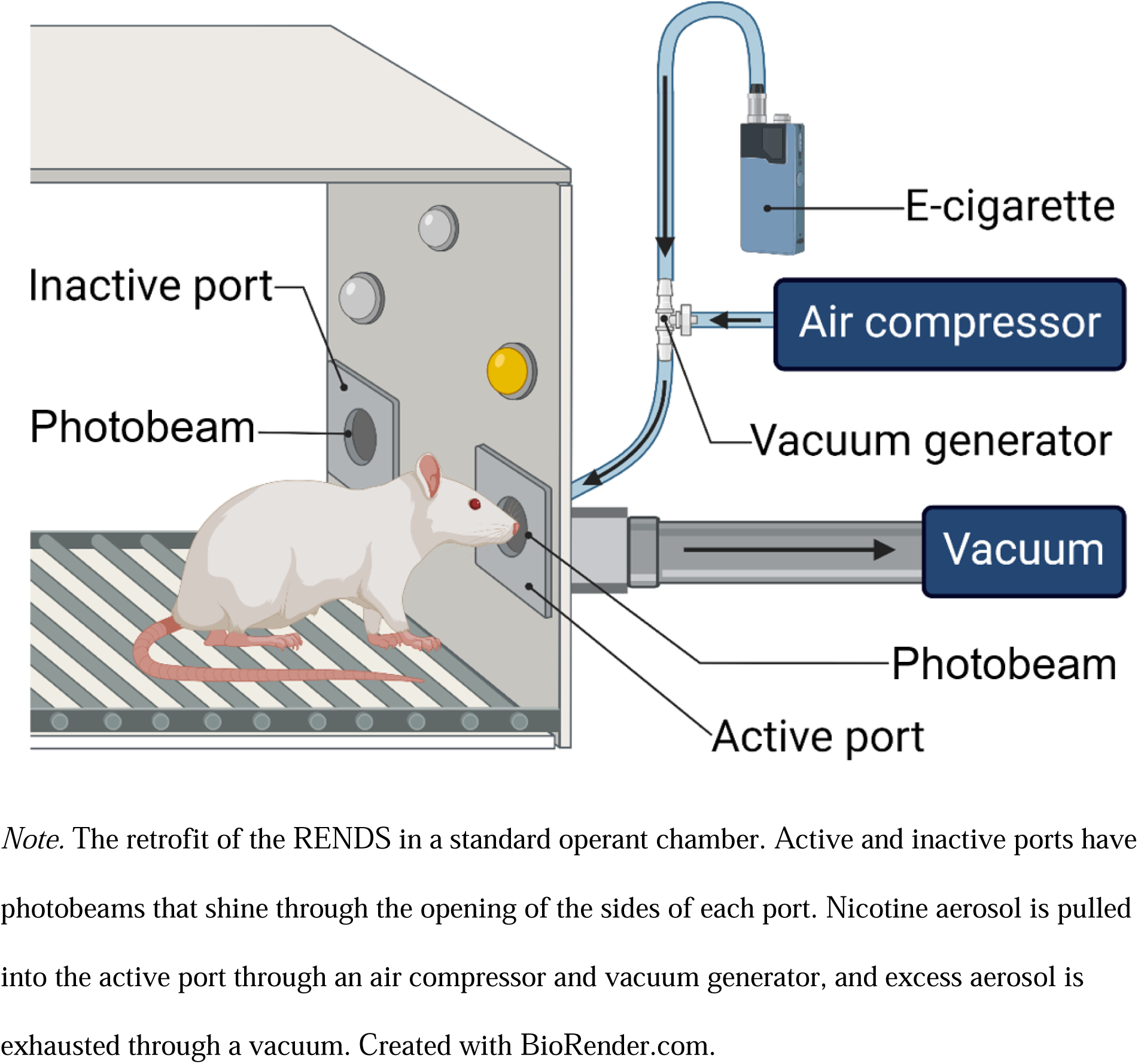
The RENDS Diagram.

At the top of the active nose port in each chamber, silicon adapters attach the nose port to the e-cigarette and air circulatory system. Air comes into the system from an air compressor (Husky 20 Gallon Vertical Electronic-Powered Silent Air Compressor 3332013) through heat-resistant pneumatic tubing (Tailonz 6mm OD 4mm ID Polyurethane PU Air Hose Pipe Tube)^2^. An air regulator (CNBTR Multicolor Acrylic 3-30LPM LZQ-7 Oxygen Air Gas Flowmeter with Control Valve) allows adjustment of the air pressure to ensure the nicotine aerosol does not exit the nose port. From the air regulator, the pneumatic tubing extends to a vacuum generator (193480, Festo) and connects the top of the e-cigarette to the top of the nose port. A custom 3D printed e-cigarette holder (Figure 2),^3^ designed in PrusaSlicer 2.9.0, and printed in polylactic acid (PLA; Hatchbox, 1.75 MM filament) on a Prusa i3 printer, is mounted to the inner wall of the sound-attenuating cubicles approximately 25.4 cm (10 in.) from the outer chamber floor. The e-cigarette is activated by a solenoid (Relay Specialties 53761-82) that manually extends to press the ‘on’ button of the e-cigarette^4^. Tubing (Cen-Tec Systems Quick Click Hose 1.25 in. diameter) attached to the end of the nose ports, secured with a stainless steel adjustable clamp (IDEAL-TRIDON 3/4-in to 1-3/4-in dia Stainless Steel Adjustable Clamp), extends to a custom 3D-printed vacuum tubing splitter, printed in PLA, which connects to the same type of tubing coming from the vacuum (Craftsman 4 gal Corded Wet/Dry Vacuum 120 V 5 HP)^5^. Air is exhausted through the vacuum, which is located in a fume hood or underneath a building air exhaust as was determined adequate by the university environmental safety department.

**Figure 2.**
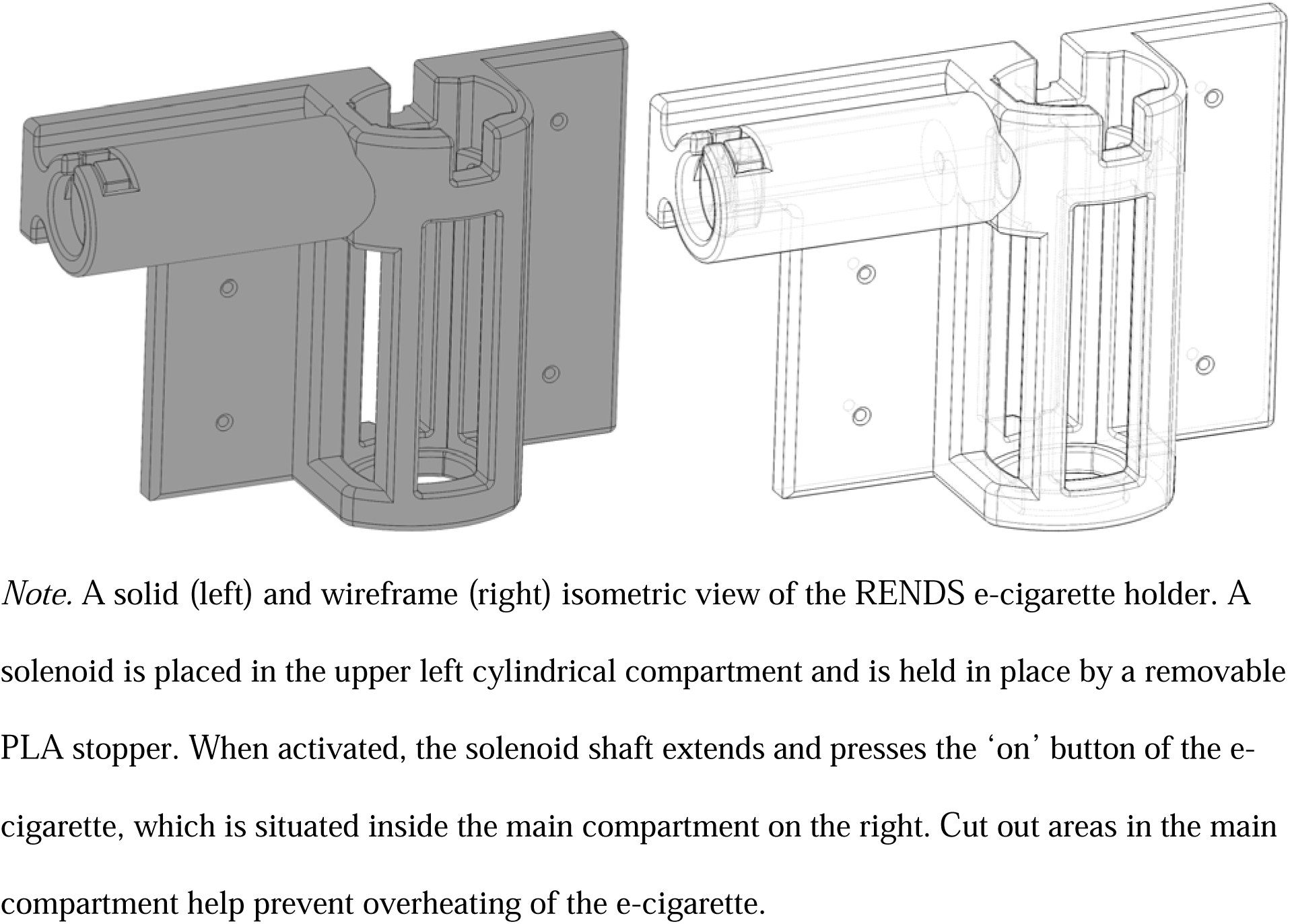
The RENDS E-Cigarette Holder.

The specially designed 3D-printed nose port^6^ (Figure 3), designed in Chitubox and printed in resin (Anycubic UV Photopolymer Resin for LCD 3D Printing, clear) on an Elegoo Saturn 8k printer, has two openings for consequence deliveries and can detect snout entry through an inset photo beam. The photobeam shines across the opening through recesses on each side of the port. Photobeams are constructed with IR break beam sensors (Electro Maker, 2167), perforated bread boards (Digikey, 1568-1652-ND), 220 OHM resistors (Virlros, VIPL080), 20-gauge speaker wire (Amazon, B09SH48KGK), and female and male 3-pin Molex connectors (Vetco Electronics, 61-403)^7^. Nicotine aerosol is delivered through a small opening in the top of the port, and then exhausted through four openings on the face of the port. In addition to the delivery of the e-cigarette aerosol, the port has a liquid reservoir to receive sucrose delivery during pre-training or other uses (e.g., ethanol delivery). The liquid reservoir is filled through a ball pump needle (SHIBUN steel air inflation needle pin A8N3) attached to heat resistant Tygon® tubing (S3 Non-DEHP 1/16”IDx3/10”OD). The needle is secured in the opening of the reservoir with industrial-strength adhesive (Eclectic E6000 PLUS). The tubing extends from the port and attaches to a 60 mL liquid syringe (BP). We deliver liquid into the reservoir using a syringe pump (PHM-200, Med Associates, St. Albans, VT), but in the absence of a pump, delivery could be achieved by manual depression of the syringe plunger.

**Figure 3.**
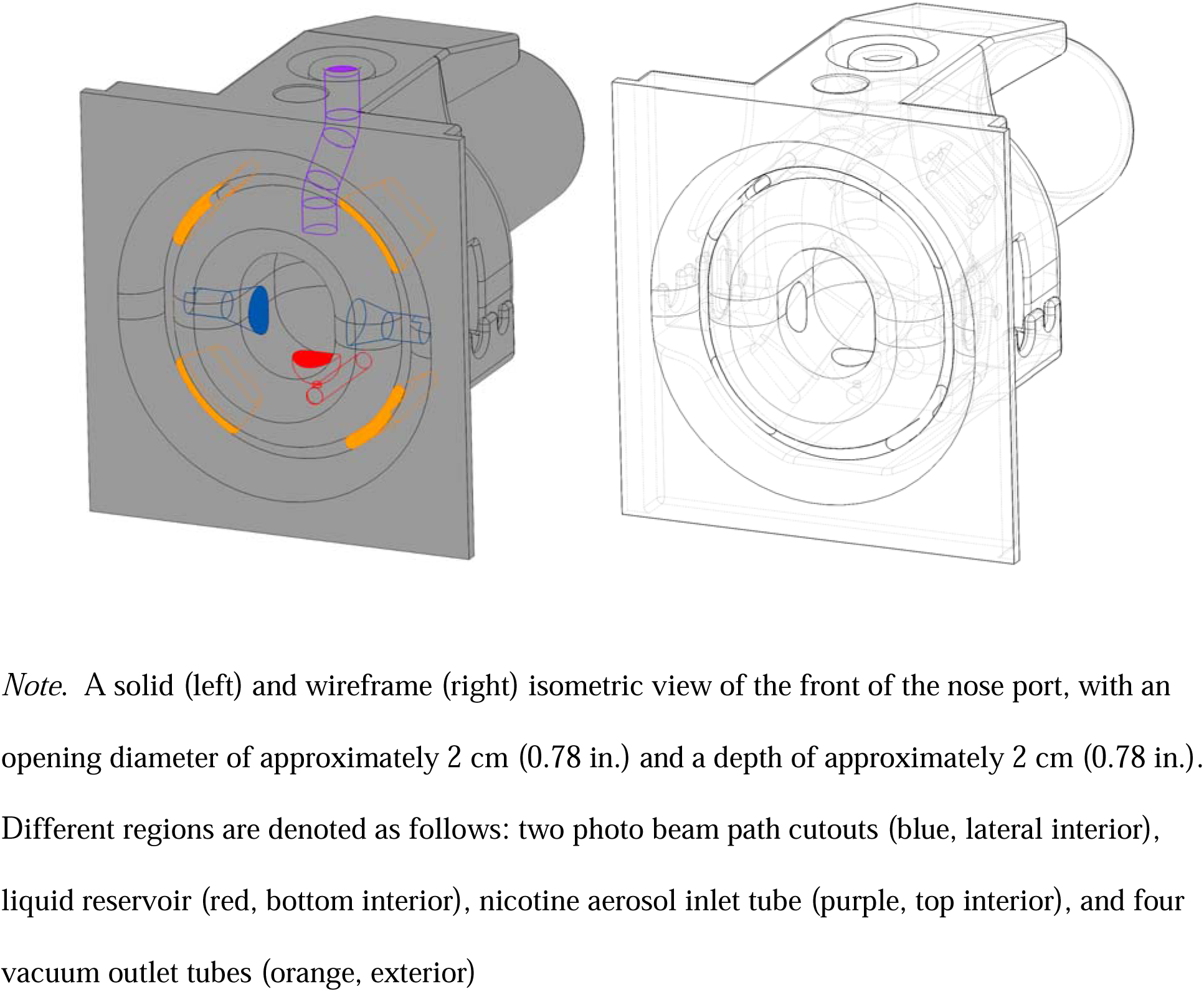
RENDS Nose Port.

##### E-Cigarette and Nicotine Solutions

We chose an e-cigarette (VooPoo Drag X mod pod; voopoo.com) and e-liquid to model current e-cigarettes and e-liquids favored by adolescents and young adults (Birdsey et al., 2023; Wang et al., 2020). Specifically, we used nicotine salt (nicotine hydrogen tartrate, SML1236; Millipore Sigma), which is easier to inhale than the harsher free-base form (Leventhal et al., 2020, 2021). E-cigarettes were set to 12 W and 1.2Ω, as recommended for salt solutions (e.g., Rubery, 2022). Mesh coils were replaced every five days, which at the stable level of self-administration of the present study amounts to after roughly 750 2-s puffs. We suspended the nicotine at 5 mg/mL with 5% mango flavoring (vaporvapes.com) in a base of 50/50 propylene glycol/vegetable glycerin (PG/VG; vaporvapes.com). This dose maintained substantial self-administration behavior in prior pilot work. We verified that the flavoring and PG/VG base had no detectable nicotine via gas chromatography/mass spectrometry (GC/MS) with a limit of detection of 0.015 mg/mL.

##### Validation

We used several pieces of apparatus and materials to validate the operation and function of the RENDS. To verify the RENDS contained the flavored nicotine aerosol in the nose port, we assessed air quality using a relatively low-cost and readily available monitor, the Zen (PurpleAir). The monitor was placed on the grate floor, equidistant from all sides of the chamber. The Zen has two sensors, and PM2.5 levels are reported as the average of the measurements from each sensor (i.e., the sum of PM2.5 levels for both sensors divided by two).

To verify that rats were ingesting meaningful levels of nicotine, blood serum was tested for the major nicotine metabolite cotinine. Nicotine subjects were anesthetized via isoflurane during blood serum collection using a VetEquip V-1 tabletop system with a scavenging cube (item no. 931600), a 2L vented induction chamber (942102), and a CX-R circuit nose cone (921616). A F/AIR charcoal filter canister (Thermo Fisher Scientific catalog no. NC9989597) attached to the scavenging cube removed excess anesthetic gas during the procedure. The induction chamber and nose cone received anesthetic gas from the vaporizer (VAS incorporated, serial no. 104417) with a refillable isoflurane adapter and an oxygen tank which was regulated by a VetEquip O2 flow meter and Pro-star oxygen regulator gauge (model no. G250-150-540). Isoflurane gas ranged from 1-5% as needed to maintain anesthesia. To analyze the obtained blood serum for cotinine, we used a Calbiotech ELISA Assay (CO096D-100) with a standard range of 0-100 ng/mL and analytical sensitivity of 1 ng/mL on a Bio Tek ELx808 microplate reader according to the included ELISA kit standard protocol. Values outside the range of the standards were calculated using the equation from the standards curve with the minimum and maximum standard values.

To assess spontaneous somatic withdrawal symptoms, rats were placed individually in a 23 cm wide x 28 cm tall glass cylinder (Tabletops Unlimited 2L Glass Apothecary Jar). A sheet of plexiglass with a 5.08 cm (2 in.) diameter ventilation hole in the center, placed on top of the cylinder and secured by a 2.27 kg (5 lb) weight, served to keep rats inside. An iPad Pro (model no. ML0G2LL/A) camera, located on a tripod 41.91 cm (16.5 in) away from the cylinder, was used to record withdrawal assessments.

### Procedure

#### RENDS Apparatus Assessment

We measured PM2.5 particulate matter in three separate conditions in the closed operant chamber. In all conditions, air quality was assessed for 10 mins. We first measured air quality in the chamber without the e-cigarette or vacuum system operating (room air). This assessment provided a baseline measure of air quality. For the following two conditions, a 2-s e-cigarette aerosol presentation occurred each minute. In the first of these conditions, the vacuum system was on, and in the second of these conditions, the vacuum system was off. Between each condition, the air quality monitor was removed from the chamber for 10 mins to allow the sensors to re-adjust before the next condition. We conducted this same assessment (baseline room air, puffs with vacuum system on, puffs with vacuum system off) on six consecutive days at approximately the same time. Air quality measurements are reported as mean levels of PM2.5 across the 10-min tests for each assessment.

#### Self-Administration

##### Behavior

The RENDS system produces a variety of sounds due to the operation of the vacuum and the periodic refilling of the air compressor. Therefore, nicotine subjects were first acclimated to the RENDS by placing their home cages in a hallway adjacent to the experimental room for 1 hr with the RENDS system running.

The apparatus was tested prior to each daily experimental session. Sessions began with a 1-min chamber blackout, during which nose pokes had no programmed consequences and visual stimuli were off. Sessions ended after a maximum of 85 mins. Nose pokes with a duration of at least 20 ms were counted except as noted during pre-training.

###### Pre-Training

Nicotine subjects underwent pre-training with a 10% (w/v) sucrose solution to hold their snout in the active nose port. All sessions during pre-training ended after 40 0.09-mL sucrose deliveries (2.84 s activation of the syringe pump) into the liquid reservoir on a Variable Time (VT) 90 s schedule using a Fleshler and Hoffman (1962) distribution. During the two sessions of magazine training, the stimulus light over the port was lit during the sucrose delivery. During the one session of autoshaping (Brown & Jenkins, 1968), the house light and the stimulus light above the nose port were lit for 10 s, followed by response-independent sucrose presentation. Any nose pokes during stimulus presentations terminated the trial and produced the immediate delivery of sucrose.

Next, the reinforcement schedule requirements were gradually increased, and rats were trained to hold their head in the port, across fifteen sessions. These schedules started with a Fixed Ratio (FR) 1, in which each response produced sucrose, then progressed through variable ratio schedules, in which the sucrose was delivered after an unpredictable number of responses with a certain mean (Ferster & Skinner, 1957) starting with a Variable Ratio (VR) 1.5 and moving ultimately to a VR3. After two sessions in which all available sucrose deliveries were obtained within the maximum session time, the required duration of the nose poke was gradually increased both within and across sessions to 2 s to ensure the rats could keep their snouts in the port for the duration of the eventual nicotine aerosol presentation. The VR3 schedule began once rats reliably earned all sucrose deliveries with the 2-s nose-poke requirement. Once the VR3 schedule began, the 2-s nose-poke requirement was removed and the minimum duration for a nose poke to be recorded was again 20 ms to ensure that rats could completely control the amount of nicotine ingested.

Throughout pre-training and the rest of the experiment, at the start of each trial, the house light and stimulus lights illuminated, and after the schedule requirement was met, the stimulus lights extinguished, and the consequence (sucrose or aerosol) was delivered; the house light turned off at the end of consequence delivery. There was a 30-s inter-trial interval (ITI), during which all lights were off, and nose pokes had no consequences, after each consequence delivery. The procedure is modeled on that used in IV nicotine self-administration (e.g., Pittenger et al., 2016).

###### Experimental Conditions

Following pre-training, the four nicotine subjects completed five experimental conditions to examine nicotine aerosol self-administration. Here, we describe and present data from three of these conditions: VR2 baseline, extinction, and VR2 return to baseline^8^. During self-administration, flavored nicotine aerosol was presented for 2 s in the nose port on a VR2 schedule with all details as in pre-training. Sessions ended after 75 deliveries of nicotine aerosol or 85 min. This condition was in effect for 50 sessions. For the extinction condition, the RENDS system and all chamber stimuli were off, and nose pokes had no scheduled consequences (i.e., nicotine aerosol was not available). Nose pokes in either port continued to be recorded. Extinction was in effect for 10 sessions. During the VR2 return to baseline period, which was identical to the VR2 baseline condition, nicotine aerosol was again available for active nose pokes; the VR2 return to baseline was in effect for 25 sessions.

###### Blood Levels of Cotinine

We analyzed blood serum samples for the major nicotine metabolite cotinine by ELISA assay. Blood serum samples were collected under anesthesia 20 min post experimental sessions via lateral saphenous vein puncture (Van Herck et al., 2001)^9^. Estimates for cotinine levels for e-cigarette users range from 100-300 ng/mL or higher, with an average around 200 ng/mL (Rapp et al., 2020; Scherer et al., 2022; Zhang et al., 2024). We obtained blood samples after session 20 and 43 (M12 & F22) and 21 and 44 (M11 & F21) of VR2 baseline, and after session 10 of extinction (F21 & F22). Males did not experience blood draws during extinction as they had adverse reactions to the anesthesia.

###### Spontaneous Somatic Withdrawal

We based the spontaneous somatic nicotine withdrawal symptom assessment on a combination of previous procedures (Besheer & Bevins, 2003; Frie et al., 2023; Malin et al., 1992). One day before withdrawal assessments, nicotine and control subjects were acclimated to the observation cylinders for 10 min. During the withdrawal assessment, nicotine rats were placed in the observation cylinders and recorded for 10 min directly before a self-administration session, approximately 22 hr after the last session. This assessment period allowed for 1) data collection during the longest duration of nicotine abstinence between sessions and 2) an optimal opportunity to observe somatic symptoms of spontaneous withdrawal at their peak following the termination of nicotine consumption in rats (approximately 17 to 23 hr; Malin et al., 1996, 1998). Control rats experienced the same assessment as the nicotine rats at approximately the same time of day. Withdrawal assessments occurred roughly two months after the last condition depicted in Figure 6, following continued daily sessions of self-administration.

After withdrawal assessments were recorded, each video was assigned to two individual coders for scoring. Teeth chattering, writhes or gasps, shakes, piloerection, yawns, dyspnea, and ptosis were all coded as target somatic symptoms of withdrawal (Frie et al., 2023; Malin et al., 1992). We developed a code book featuring the operational definitions and examples of each symptom for review before scoring, which can be found on OSF. We used partial interval scoring (i.e., noting either yes or no for the occurrence of each symptom) during one-min intervals throughout the 10 min assessment period. Interobserver agreement (IOA) was calculated by dividing minute interval agreements by the sum of agreements and disagreements. Final withdrawal scores were found by averaging the two ratings (Frie et al., 2023).

### Data Analysis

Most papers published in *JEAB* follow an analysis strategy of presenting individual subject data for visual inspection as well as performing inferential statistics (Kyonka et al., 2019), and we followed this convention. For all analyses, we relied mainly on visual inspection of individual subject data (Perone, 1991), but, when possible, we also included inferential tests to further delineate findings and to provide estimates of effect sizes^10^. Data were collapsed across sex for each analysis due to the small sample size. Prior to completing inferential statistics, we conducted Shapiro-Wilk tests for normality (Shapiro & Wilk, 1965). All analyses and data visualizations were conducted in GraphPad Prism (version 10 for Mac OS, Boston, MA). Prism files are found on OSF.

Our first analyses focused on the apparatus assessment of PM2.5 levels across conditions of room air, puffs with the vacuum system on, and puffs with the vacuum system off. All data except for room air passed normality testing; room air, *W* = 0.50, *p* < .001. Therefore, for the apparatus assessment, we compared PM2.5 levels in the chamber with room air and puffs with the vacuum system on using a nonparametric Wilcoxon matched pairs signed rank test. In comparing PM2.5 levels for puffs with the vacuum system on and off, a paired-samples *t* test was performed. We report Cohen’s *d* effect sizes with conventions described as follows: small effect = .2, moderate effect = .5, and large effect = .8 or greater (Cohen, 1988); however, these conventions are generally noted as arbitrary and should be treated as guidelines (Lakens, 2013).

Our second set of analyses focused on self-administration assessments. Specifically, we examined nose pokes in active and inactive ports across baseline VR2 sessions, extinction, and return to baseline VR2 conditions, collapsed across sex. We note any observed sex differences via visual analysis of the disaggregated data (see Odum et al., 2024). We first show representative cumulative records for each subject on the last session of each condition. We then show the total number of active and inactive nose pokes for the last six sessions of each condition for individual subjects. With the exception of active nose-poke responses during extinction, all data from the last six sessions for each condition were non-normally distributed: baseline VR2 active nose pokes (W = .74, *p* < .0001); baseline VR2 inactive nose pokes (W = .84, *p* =.001); extinction active nose pokes (W = .98, *p* =.85); extinction inactive nose pokes (W = .77, *p* <.001); return to VR2 baseline active nose pokes (W = .68, *p* <.001); return to baseline VR2 inactive nose pokes (W = .80, *p* =.001). Therefore, we first transformed data by ranks and conducted a 2 (nose pokes: active vs. inactive) by 3 (condition: baseline VR2, extinction, return to baseline VR2) nonparametric-approximate mixed ANOVA (a two-way nonparametric test for repeated measures comparable to a Friedman’s test with interactions; Conover & Iman, 1981; Friedman, 1937)^11^. Bonferroni multiple comparison tests were used to examine more specific differences in the ranked data^12^. We report effect sizes as η^2^ with the following conventions: small effect = 0.01, medium effect = .06, and large effect =.14 or greater (Adams & Conway, 2014; Cohen & Cohen, 1983).

For the next analyses, we examined the cotinine and withdrawal data. Because the cotinine data were non-normally distributed (W = .82, *p* = .032), we computed a Spearman’s rank-order correlation between the number of puffs earned for the VR2 baseline and extinction sessions and resulting cotinine levels for those sessions We report Spearman’s ⍰ as the effect. Finally, we compared withdrawal scores, collapsed across sex, for the nicotine and control size subjects using an independent sample *t*-test. We report Cohen’s *d* for effect size, using the conventions we described above in the *t*-test analyses for air quality; effect size conventions for ⍰ are the same as Cohen’s *d*.

## Results

### Apparatus Assessment

The level of PM2.5 particles in the chamber air with puffs with the vacuum system on was similar to room air level with no puffs (see Figure 4). Specifically, in the room air condition, PM2.5 was very low (*M* = 0.17 µg/m^3^, *SD* = 0.41). The PM2.5 level for puffs with the vacuum system on (*M* = 2.58 µg/m^3^, *SD* = 2.11) did not differ from room air with no puffs (*n* =12, *W* = 15, *p* = .063). The means for the room air with no puffs and vacuum system on with puffs were both well below the cut-off for the highest standard for air quality (see Figure 4 note). Finally, in the condition with puffs with the vacuum system off, PM2.5 was extremely high (*M* = 2,631.5 µg/m^3^, *SD* = 241.6) and over 10 times the level considered hazardous (see Figure 4 note). Mean PM2.5 levels were significantly different between puffs with the vacuum system on and off (*t*(5) = 26.66, *p* < .0001).

**Figure 4.**
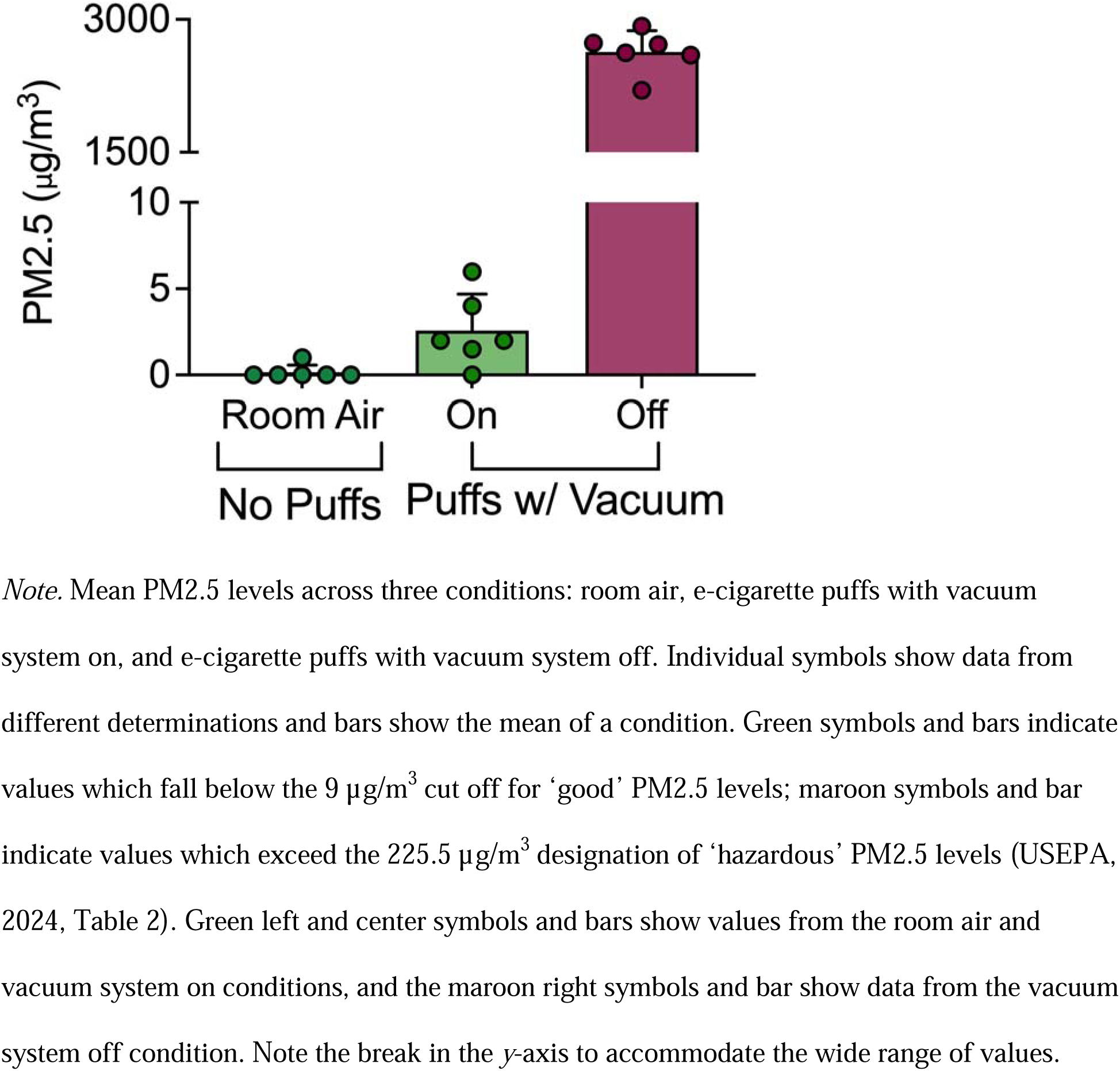
Measurements of Chamber Air Quality Across Three Conditions.

### Self-Administration

Figure 5 shows that the cumulative number of active port nose pokes during the self-administration VR2 baseline and return to VR2 baseline sessions far exceeded that of the number of inactive port nose pokes for all rats. During baseline and the return to baseline, active port nose pokes typically started 5-10 mins after the beginning of the experimental session and occurred at a high constant or increasing rate. Furthermore, when active port nose pokes no longer produced the flavored nicotine aerosol and stimuli in the extinction condition, active and inactive nose pokes occurred at a similarly low rate across the session. Male rats made more inactive nose pokes than female rats during baseline and return to baseline. The number of nose pokes in the active port, as well as the difference between nose pokes in the active and inactive ports, was relatively similar during the baseline and return to baseline sessions.

**Figure 5.**
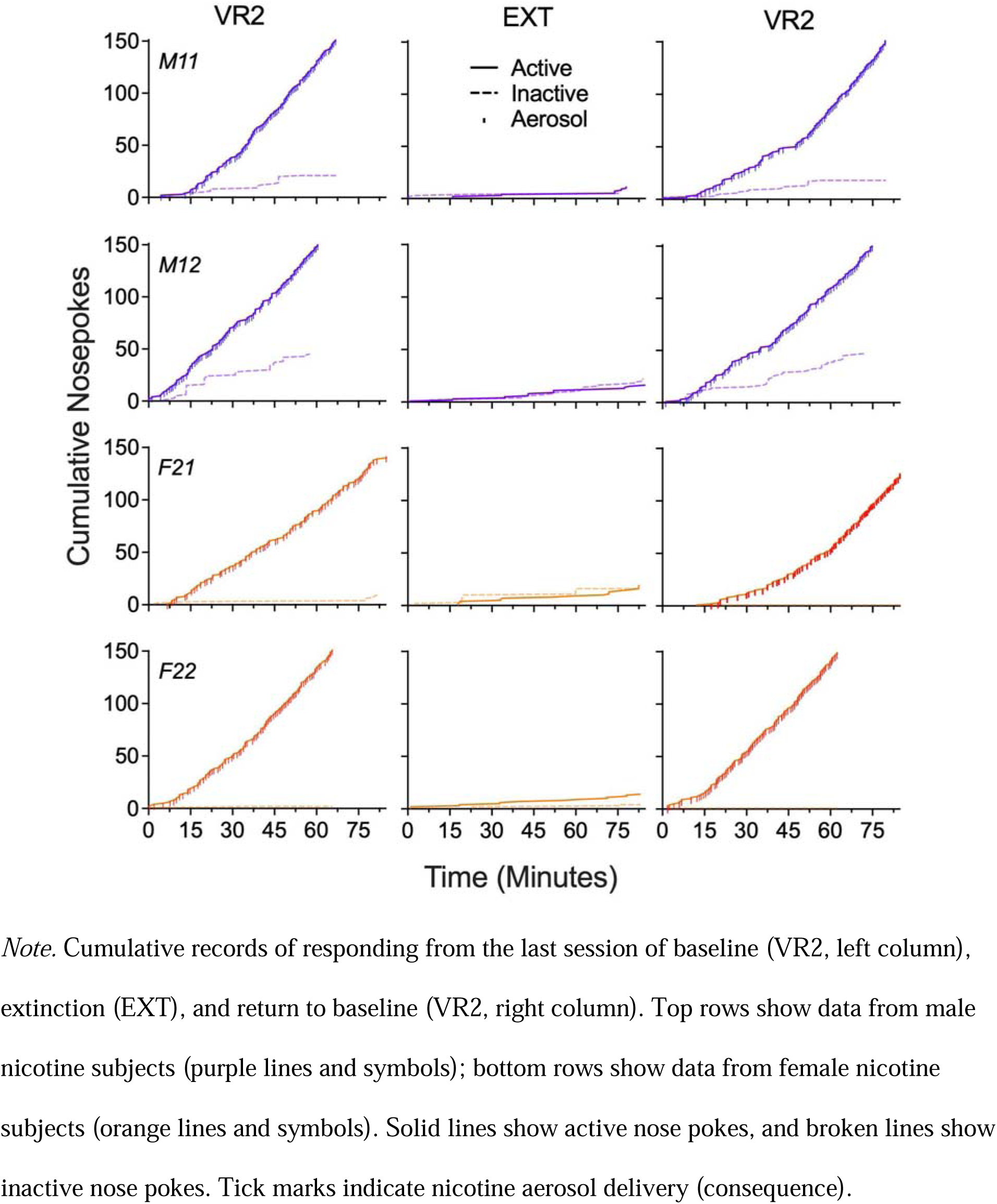
Cumulative Records for Baseline (VR2, Left Column), Extinction, and Return to Baseline (VR2, Right Column) Conditions for Male and Female Rats.

Figure 6 shows the total number of active and inactive nose pokes for the last six sessions of the self-administration VR2 baseline, extinction, and return to VR2 self-administration conditions. Nose poking differed by condition, as shown by a significant main effect of condition, *F* (2, 69) = 29.87, *p* <.0001, η^2^ = 15.24. There was also a significant main effect of nose pokes, *F* (1, 69) = 170.8, *p* <.0001, η^2^ = 37.69, showing that the total number of active nose pokes greatly exceeded that of inactive nose pokes across conditions. Additionally, there was a significant nose poke by condition interaction *F* (2, 69) = 32.25, *p* < .0001, η^2^ = 14.23, showing that active and inactive nose pokes were affected differently across conditions.

**Figure 6.**
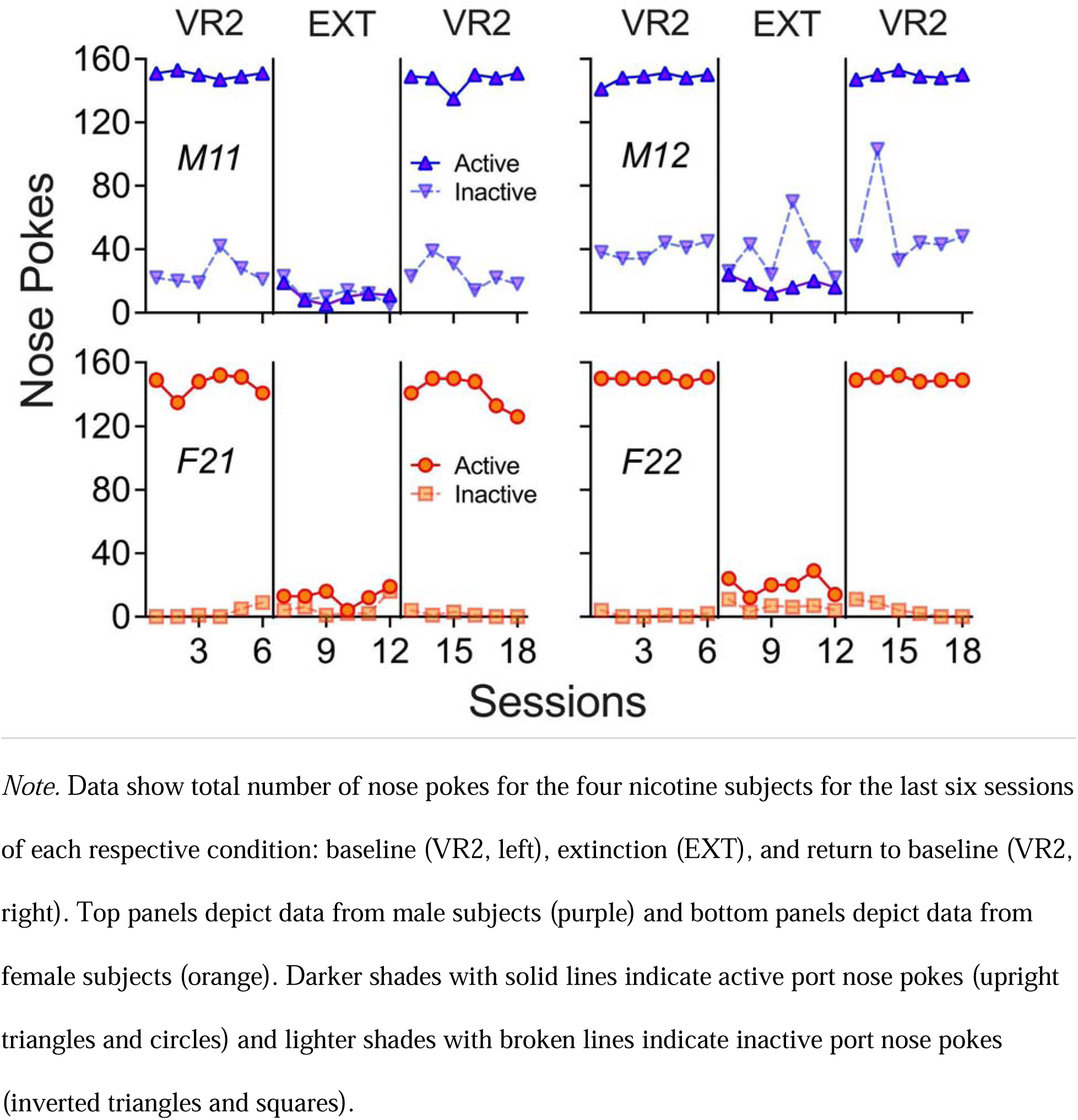
Total Nose Pokes Across the Last Six Sessions for Each Rat for Each Condition.

The total number of active nose pokes was high and consistent across the sessions of baseline and return to baseline conditions. The average across all rats was 147.65 active nose pokes, which represents a ceiling due to the limit of 75 aerosol presentations. In these sessions, there were no differences in the number of active nose pokes for male and female rats. In follow-up tests, active nose pokes (*M* = 148.5, *SD* = 4.065) were significantly greater than inactive nose pokes (*M* = 17.08, *SD* = 17.14) during the VR2 baseline (p <.0001) as well as for the VR2 return to baseline (active nose pokes, *M* = 146.8, *SD* = 6.53; inactive nose pokes, *M* = 20.63, *SD* = 24.32; *p* <.0001). In extinction, active nose poking fell drastically (*M* = 15.29, *SD* = 6.011) to 10% of the number of active nose pokes during baseline; active and inactive nose pokes (*M* = 15.29, *SD* = 16.43) did not significantly differ during extinction (*p* =. 31). Inactive nose pokes remained relatively low and unchanging across conditions, with 17.77 inactive nose pokes on average across all conditions. Male rats made more inactive nose pokes than females, with M12 in particular showing a higher and more variable number of inactive port nose pokes. The ratio of active to inactive nose pokes, averaged across baseline conditions, was 6:1 (M1), 3:1 (M2), 76:1 (F1), and 81:1 (F2); averaged across rats, the ratio was 42:1.

Figure 7 shows a strong positive correlation between the number of puffs earned and cotinine levels for nicotine subjects during the VR2 baseline (Spearman’s ⍰ = .84, *p* = .01). This relation confirms that as the number of puffs earned increased, blood levels of cotinine increased. Blood serum from nicotine subjects had high cotinine concentrations during the VR2 baseline condition. Specifically, the average cotinine concentration across males and females was 153.75 ng/mL. The range of cotinine during the VR2 baseline was 12.12 to 214.39 ng/mL (the highest level of cotinine that could be calculated with the ELISA kit). During extinction, cotinine levels for female subjects were very low (*M* = 1.63).

**Figure 7.**
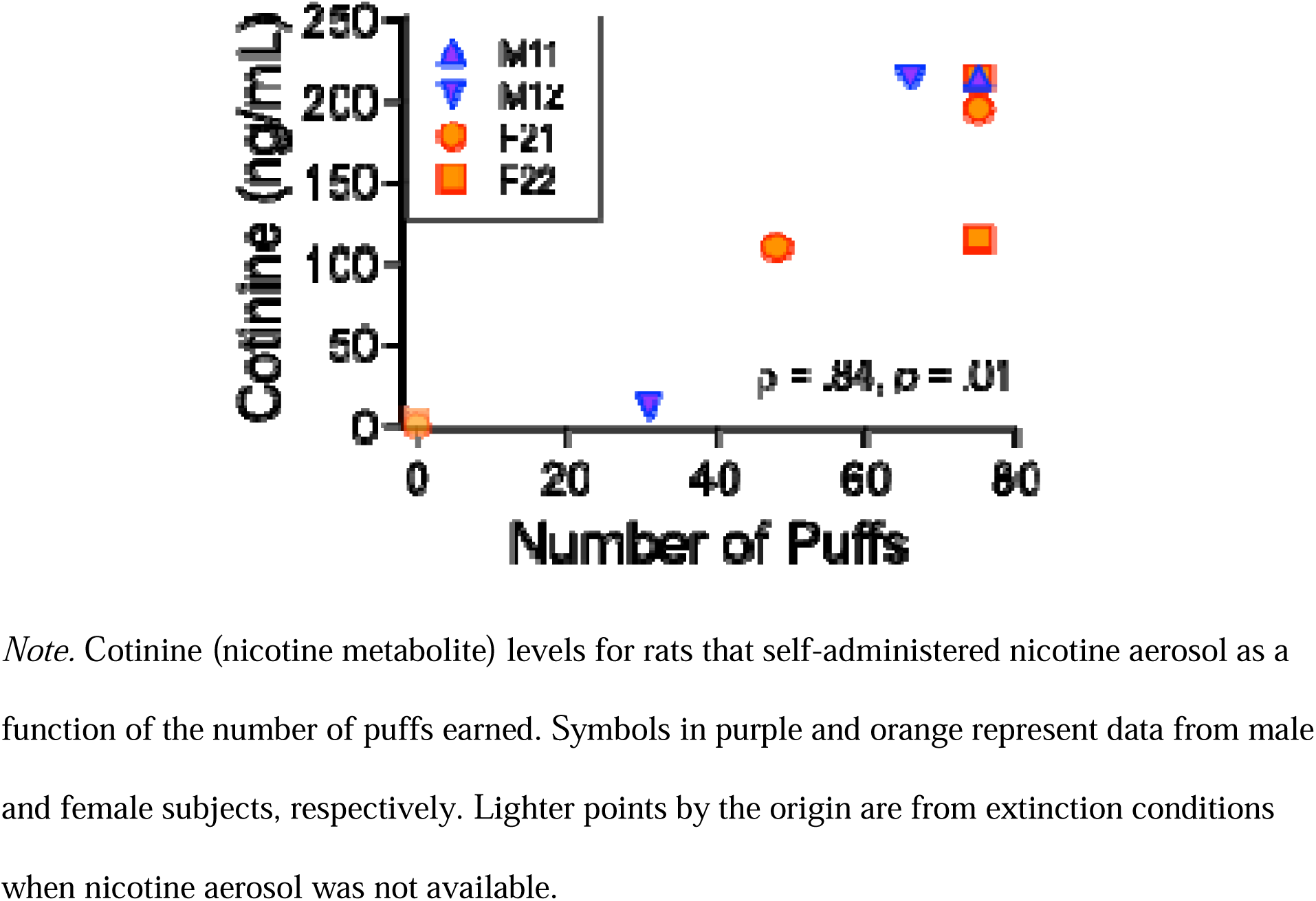
Blood Levels of Cotinine as a Function of E-Cigarette Puffs Earned.

Figure 8 shows that scores on the withdrawal scale were substantially higher for nicotine rats than control rats. Specifically, mean scores were significantly different between nicotine (*M* = 31.13, *SD* = 3.01) and control subjects (*M* = 7.00, *SD* = 1.41; *t*(4) = 10.31, p < .0005). The difference in withdrawal scores between nicotine and control subjects was large, *d =* 10.26. Female nicotine rats had somewhat lower withdrawal scores than males. Mean IOA across all symptoms was acceptable at 80.36%. Withdrawal scores for specific symptoms for all subjects can be found on OSF.

**Figure 8.**
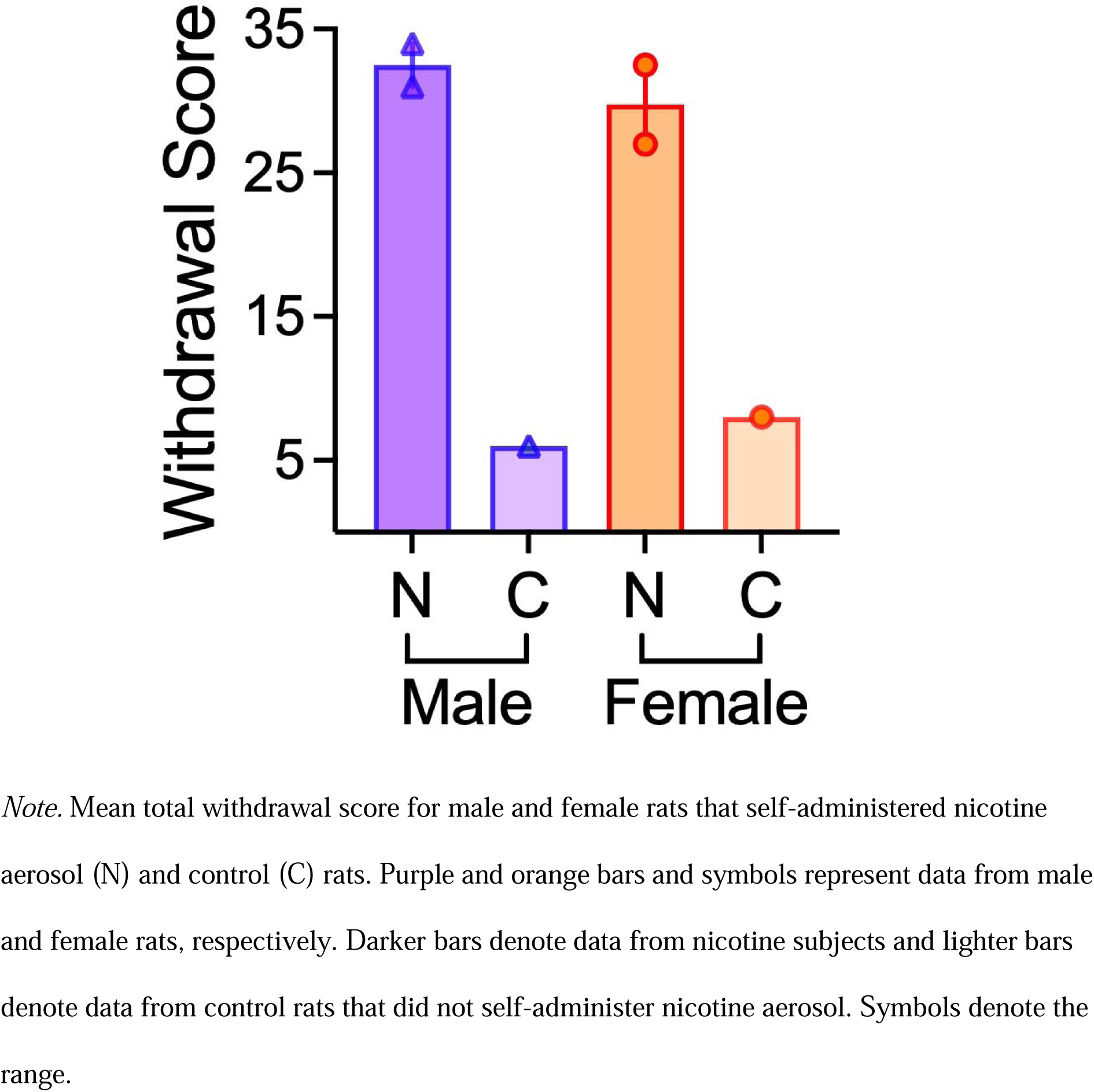
Overall Occurrence of Spontaneous Somatic Withdrawal Symptoms.

## Discussion

We described and validated the RENDS, a novel preclinical system for nicotine aerosol self-administration, across two main assessments: apparatus operation and self-administration. Our goal was to advance preclinical research on e-cigarette use with a low-cost non-proprietary laboratory animal model with high face validity. Specifically, we found that the system sequestered nicotine aerosol in the nose port, shown by similar levels of PM2.5 particulates between the air in the operant chamber (room air) without any e-cigarette puffs and the chamber air with puffs and the vacuum system in operation.^13^ We also found that rats reliably self-administered nicotine aerosol through the nose port, demonstrated by a high number of active nose pokes, cotinine levels, and withdrawal symptoms. Below we describe and discuss these findings in turn.

To determine whether RENDS functions as intended, by sequestering nicotine aerosol in the nose port, we evaluated the level of PM2.5, extremely small particulates produced by e-cigarettes (e.g., Li et al., 2020; Shearston et al., 2023). We found no difference between the levels of PM2.5 for baseline room air compared to PM2.5 levels when the vacuum system was on. This result verifies that when the RENDS system is in operation, e-cigarette PM2.5 particles do not enter the chamber at a rate above that found in room air. Thus, to consume nicotine with the RENDS, rats must access the aerosol inside the nose port. We also found, unsurprisingly, that when the vacuum system was not in operation and the aerosol filled the chamber, as in vapor chamber models, PM2.5 levels were significantly higher. Together, these results highlight that RENDS provides access to inhaled nicotine aerosol through snout-only contact with the port, which not only increases face validity but eliminates whole-body contamination issues with vapor chamber models (Chellian et al., 2023; Kogel et al., 2021; Lallai et al., 2021; Lucci et al., 2020). Furthermore, it documents that PM2.5 levels in a closed chamber with one 2-s puff per min for 10 min reach levels over 100 times higher than considered hazardous by the USEPA (2024).

Our second form of validation of the RENDS focused on self-administration. In the behavioral assessment, we examined nose poking in the active port for flavored nicotine aerosol when it was available (VR2 baseline), when access was no longer available (extinction), and when nicotine aerosol was again available (VR2 return to baseline). This rigorous reversal design showed that overall, rats robustly self-administered nicotine aerosol when available (average 147.65 active nose pokes per session), but showed minimal active nose poking when nicotine aerosol and associated cues were not available (average 15.29 per session during extinction). Furthermore, nose pokes occurred at a far higher rate in the active port, which produced nicotine aerosol, than the inactive port, which did not produce nicotine aerosol (average ratio of 42:1 across rats and baseline conditions). These findings are clear both in cumulative records of nose poking across time in the last session of each condition (Figure 5) and in total nose pokes per session across the last 6 sessions of each condition (Figure 6). Furthermore, these findings demonstrate much stronger behavioral control by the response contingencies compared to the vapor chamber full body exposure method (e.g., Cooper et al., 2024; Marusich & Palmatier, 2023; Smith et al., 2020).

Although prior research has found some evidence of discrimination between active and inactive responses with nicotine aerosol self-administration in the vapor chamber method, our study is the first to show robust discrimination. In previous work, response rates were generally low on both the active and inactive operanda (e.g., Cooper et al., 2024; Marusich & Palmatier, 2023; Smith et al., 2020). For example, Smith et al. (2020) found 20 or fewer active lever presses across 1 hr sessions, and a similar or higher number of inactive lever presses. In a study with e-liquid composition similar to that in the present research, but with mice in 2-hr sessions in vapor chambers, Cooper and colleagues (2024) found less than 15 active nose pokes (7.5/hr) with green-apple flavored 6 mg/mL nicotine salt. Furthermore, mice first had involuntary aerosol pre-exposure to induce nicotine dependence and continued to receive response-independent puffs during sessions if there was a long pause in nose poking. Additionally, in these studies a large percentage of mice (> 30%, Cooper et al., 2024; > 75%, Henderson et al., 2024) were excluded from the study for maintaining a less than 2:1 ratio of active:inactive nose pokes. In the present study, rats averaged over 105 active pokes/hr and a ratio of active:inactive pokes of over 40:1 with no prior nicotine exposure or response-independent puffs, and 100% of subjects were included in the study and vigorously self-administered nicotine aerosol.

There may be several explanations for the difference in the overall number of active responses as well as better discrimination between active and inactive responses shown with RENDS compared to the vapor chamber model. First, we used the salt form of nicotine and most other prior studies used the free-base form (e.g., Smith et al., 2020; Marusich & Palmatier, 2023); however, this difference seems an unlikely explanation because Cooper et al. (2024) used the salt form and still found low levels of active responses. Furthermore, we used flavored nicotine aerosol and most other studies have used unflavored; however, this difference also seems an unlikely reason because studies using flavored nicotine with the vapor chamber method have also found a low number of active responses and poor discrimination (e.g., Cooper et al., 2024; Marusich & Palmatier, 2023). Additionally, poor discrimination and low active response rates have been found using the vapor chamber method with both rats and mice (e.g., Cooper et al., 2024; Smith et al., 2020), so the species seems an unlikely cause for the differences as well.

There are more likely reasons for the robust active responses with the RENDS compared to the vapor chamber method. First, compared to the vapor chamber, there is greater temporal and spatial contiguity between the active response and aerosol delivery. With RENDS, the location of the response that causes the aerosol delivery is in the same place as the source of the aerosol delivery, so the aerosol delivery is highly contiguous with the nose poke which produced it. In contrast, in the vapor chamber method, the response that produces the aerosol is removed from where the aerosol is delivered, and there is a delay from aerosol initiation to when it reaches the rodent. In general, delays between when a response occurs and a consequence is delivered weaken the response and reduce the discriminability of the contingent relation (Lattal, 1975, 1995). Indeed, Henderson et al. (2024) found that mice that did not stand close to where the nicotine aerosol entered the chamber did not acquire nose poking. A second reason for less robust self-administration in the vapor chamber technique is that the manner and length of the nicotine aerosol presentation may be aversive. The aerosol in the chamber may plausibly irritate the eyes, and the effects of higher nicotine doses are aversive (e.g., Koffarnus & Winger, 2015; Shoaib & Stolerman, 1995). The presentation of the aerosol lasts up to 100 s and rats may modify their breathing to intake less aerosol (Lallai et al., 2021). In contrast, in the RENDS, the aerosol only contacts the snout, the puff duration (2 s) is similar to that of human e-cigarette users, and rats can discontinue inhaling at any point during puffs. An additional difference is the use of VR schedules with the RENDS, rather than fixed-ratio (FR) schedules of aerosol delivery as is commonly arranged. Variable-ratio schedules may contribute to higher rates of drug self-administration and active lever discrimination (Corley et al., 2024).

Despite the advantages of strong contiguity between the response and consequence in the RENDS, this method also makes it impossible to distinguish between nicotine seeking and nicotine consumption. Specifically, subjects engage in the same response (i.e., active port nose poking) to both earn and consume the nicotine aerosol. These goal-directed and consummatory responses could individually provide enlightening insights into the fundamental nature of nicotine aerosol self-administration. For example, rats could earn aerosol presentations by pressing a lever placed underneath or beside the port and consume the presentations by placing their snout in the port, much like with lever pressing for food-maintained behavior in which the operandum and consequence delivery are spatially separated. Investigators may want to consider this sort of method of parsing these responses in the future.

Rats who self-administered nicotine aerosol through the RENDS had high blood levels of cotinine (the major nicotine metabolite). Thus, not only were the rats engaging in robust nose poking in the active port, but they were also consuming biologically relevant levels of nicotine, similar to that obtained in preclinical models of IV nicotine self-administration (e.g., Shoaib & Stolerman, 1999) as well as with humans with daily tobacco and e-cigarette use (e.g., Rapp et al., 2020; Schroeder et al., 2014). Furthermore, there was a direct relation between the number of nicotine aerosol puffs and cotinine: the more puffs earned, the higher the cotinine level. Women using e-cigarettes have higher cotinine levels than men (e.g., Park & Choi, 2019), but we were not able to examine preclinical sex differences in cotinine with nicotine aerosol self-administration due to our small sample size and ceiling effect. Future research in this area should remedy these limitations (see Odum et al., 2024). Overall, there has been less preclinical research with females than with males for voluntary nicotine use in general (Chellian et al., 2023). However, some studies with IV nicotine have found greater self-administration in female rats than male rats (Barrett et al., *in press*; Chellian et al., 2024; Leyrer-Jackson et al., 2021; Lynch, 2009; Swalve et al., 2016).

We found spontaneous somatic symptoms of nicotine withdrawal following self-administration of nicotine aerosol with the RENDS, which to our knowledge is the first in the literature. Nicotine rats had substantially higher withdrawal scores relative to control rats. The difference in withdrawal symptoms between the nicotine and control subjects replicates the only other finding of withdrawal in the literature with self-administered nicotine aerosol to date (Smith et al., 2020), which was precipitated by the nicotine antagonist mecamylamine. One potential reason for the robust spontaneous withdrawal found here is the length of nicotine aerosol self-administration before withdrawal testing (146 daily sessions). We did not evaluate withdrawal early in the self-administration condition. It will be interesting and important to examine the development of withdrawal across acquisition and maintenance conditions of nicotine aerosol self-administration.

The long-term exposure to nicotine described in this report demonstrates that our system is particularly advantageous for chronic self-administration of nicotine aerosol. Specifically, other commonly used techniques (e.g., IV self-administration) have been challenged to evaluate nicotine self-administration for long durations due to catheter issues (Suarez et al., 2024). Catheters used in IV self-administration are difficult to retain over time due to loss of patency and growth of rodents. Catheters are also challenging to implant for smaller rodents like mice and adolescent rats (Belluzzi et al., 2005; Pogun et al., 2017). Our system, however, can maintain consistent consumption of nicotine aerosol over a long time, demonstrated by self-administration by 100% of nicotine subjects across more than 200 daily sessions. In contrast, for IV catheters, 88% patency for 46 days is considered exceptional (Suarez et al., 2024). Additionally, the RENDS could be used with smaller rodents by decreasing the size of the port and lowering placement on the front panel. Thus, the RENDS, both in design and methodology, may serve as a practical and instrumental avenue for novel longitudinal investigations of nicotine aerosol self-administration in rodents.

Although we have refined the RENDS over many prototypes, the system and methodology could still be improved. For example, one limitation of the present validation was reliance on a small convenience sample of subjects of different strains and ages. This strategy was a conscious choice to reduce animal numbers in line with the three Rs of research (replacement, reduction, and refinement; NC3Rs, N.D.). The contribution of this technical report is therefore the apparatus and general procedure, rather than specific empirical findings. Future investigations in this area should increase the number of subjects as well as include matched controls to examine whether the empirical findings reported here are replicable and generalizable. In addition, during experimental conditions, we limited the number of aerosol presentations available in a session to 75. Our reasoning was to reduce the likelihood of subjects consuming high, potentially aversive, doses of nicotine. However, the maximum number of aerosol presentations was reached in most sessions, usually well before the maximum session time, leading to a ceiling effect. Future research in this area should increase the number of available nicotine aerosol presentations within a session. Additionally, future research would be strengthened by examining self-administration of nicotine aerosol as a function of dose of nicotine. Finally, it will also be important to examine choice of a flavored PG/VG vehicle over a nicotine-containing aerosol.

Although rats robustly self-administered nicotine aerosol, unlike with IV self-administration, each consequence presentation could differ in the amount of nicotine based on how long and intensely the subjects inhaled the aerosol. Therefore, we measured blood levels of cotinine, which were on average high. Blood collection is an invasive process, and usually requires that rats be either anesthetized (as in the present study) or restrained, both of which can affect self-administration and physiological data. Therefore, an alternative method for evaluating cotinine levels would be beneficial. For example, rats can be trained to accept tail vein blood sampling by skilled personnel without restraint (NC3Rs, 2022). Additionally, cotinine can be detected in rodent urine via ELISA (e.g, Calbiotech CO096D-100), which can be collected relatively easily (Kurien et al., 2004) and could provide an alternative to blood collection.

Our methodology demonstrates the strong reinforcing function of the flavored nicotine aerosol, but we did not evaluate to what extent the flavor or nicotine were responsible for the effects. Our goal was to provide a model of human e-cigarette use, because flavoring plays a powerful role in ENDS initiation and use (e.g., Audrain-McGovern et al., 2016; Baker et al., 2021; DeVito & Krishnan-Sarin, 2018; Leventhal et al., 2019, 2020; Zare et al., 2018). However, nicotine has complex functions in maintaining behavior: it can serve as a primary reinforcer (Chaudhri et al., 2007; Donny et al., 1998), imbue associated stimuli with conditioned reinforcing function (Wilkinson & Bevins, 2008), and enhance the rewarding properties of other stimuli (Barrett & Bevins, 2012; Barrett & Odum, 2011; Bevins & Palmatier, 2004). These latter processes could enhance the rewarding value of flavor and nicotine-paired cues, potentially increasing self-administration. Both flavor and cues can also maintain self-administration in the absence of nicotine (e.g., Barrett & Odum, 2011; Cooper et al., 2023). Therefore, we are unable to delineate the separate roles of nicotine, flavoring, and cues in producing the robust self-administration found with the RENDS. Whether flavor should be included in an e-cigarette protocol will depend on the goals of the research. Future research should attempt to delineate the roles of the complex stimuli present in ENDS use now that we have established a system for doing so.

An additional consideration for investigators moving forward is the method of pre-training employed in this report. Although our procedure is modeled after those commonly used in the drug self-administration literature and nicotine aerosol (e.g., Abraham et al., 2023; Pittenger et al., 2016; Marusich & Palmatier, 2023), there are adjustments to pre-training that reduce associations between the active manipulandum and food. For example, subjects can be food trained on both response manipulanda (i.e., both the eventual active and inactive) as in Barrett and Bevins (2013). This procedure ensures that responding on the active manipulandum following pre-training is explained solely by the contingency between the manipulandum and the consequence, rather than disproportionate pre-training experience and association with food on just one manipulandum. This pre-training methodology could be implemented in our procedure to increase rigor. Specifically, because subjects would have experience working for sucrose in both nose ports, disproportionate nose poking in the active port would reflect only the contingencies between nose poking in that port and the nicotine aerosol. Because our experiment continued for many sessions, it seems unlikely that the differential pre-training affected the long-term results, but non-differential pre-training would be an improvement regardless.

There are several future directions to note given that the robust data described here have provided comprehensive validation of the RENDS. Because our system addresses many concerns of other nicotine aerosol self-administration models, the RENDS can be used to evaluate the effects of long-term (chronic) and more face valid consumption of nicotine aerosol. The RENDS could also be useful in streamlining methodologies used in nicotine aerosol self-administration research. Prior research has generated mixed results, such as varying rates of self-administration and discrimination between active and inactive manipulanda (see Marusich & Palmatier, 2023 for discussion), that could be potentially explained by differential procedures and components of nicotine aerosols. With the high experimental control and discrimination between ports we have demonstrated with the RENDS, manipulation of other procedural details could generate valuable insights into the intricacies of self-administration of nicotine aerosol in preclinical analogs and clarify the mixed findings of past research.

In sum, the RENDS provides a validated low-cost system for snout-only e-cigarette aerosol inhalation in rodents, continuing prior efforts to make behavioral research more accessible (e.g., Aguiar et al., 2020; Buscher et al., 2020; Escobar et al., 2022; Escobar & Pérez-Herrera, 2015; Gurley et al., 2019; Rizzi et al., 2016). The plans for the custom-created 3D-printed components of the RENDS are freely available on OSF. Our goal is to further open the possibilities of preclinical research into ENDS with robust methods to promote regulation, prevention, and cessation strategies. It has become increasingly crucial to develop models like the RENDS to investigate nicotine aerosol self-administration, as nicotine consumption is on the rise, even as the long-term impacts and harms associated with ENDS use are not completely understood (Baldassarri, 2020; Shehata et al., 2023; Vu et al., 2024). The RENDS also enables research into the basic mechanisms that make this relatively new and pernicious form of nicotine use so appealing. Thus, we hope to foster continued research on this complex drug with widespread use dating back over 12,000 years (Duke et al., 2022).

## Acknowledgements

The authors express their gratitude to the many individuals who contributed to this project. The authors acknowledge prior graduate students Ryan Becker, Devanio Cousins, Annie Galizio, D. Perez, and Caroline Towse for their collaboration and support on earlier versions of this project. We thank Spencer Mathias for his help designing the original nose port. We also thank Megan Raddatz for helping with the blood sampling process. The authors thank Josephine Hannah for assistance in withdrawal video coding, as well as Avonlea Richards and Dillon Marstella as exceptional paid research assistants and supervisors. The authors recognize the dedicated research assistants Erin McQuillen, Kelsey Summarell, Bailey Herbert, Brooke Piippo, and Jesslyn Hoffman, whose commitment and hard work were invaluable. Additionally, the authors appreciate the contributions of research assistants who worked with us for one semester: Kaylana Brown, Callissa Candalot, Katherine Flores-Cabrera, Elisabeth Himelberger, Lexie Myers, Brandi Smith, and Ryan Thrasher. Finally, we would like to acknowledge our research subjects for their contributions to our science.

## Funding Statement

This research was supported by a Cutting-Edge Basic Research Award (CEBRA) from the National Institute on Drug Abuse (RD21 DA053818) and by a SPARC (Seed Program to Advance Research Collaborations) grant from USU.

## Footnotes

1 We retrofitted the RENDS to our existing Med Associates operant chambers, but with slight modifications it could be applied to any similar operant chamber.

2 One air compressor and vacuum can support up to six chambers fitted with RENDS.

3 The holder is specific to a VooPoo Drag X (voopoo.com) but could be readily modified for other e-cigarettes.

4 In our experience, activation longer than 2 s and more often than twice per min causes overheating of the e-cigarette.

5 E-liquid condenses in the vacuum filter, so regular cleaning of the vacuum and filter, while wearing Personal Protective Equipment (PPE) to avoid nicotine exposure, is needed.

6 The shape of the port and the resin material makes chewing less likely.

7 The recesses do not open into the port, and the photo beam instead shines through the resin, because contact with the nicotine aerosol destroys the sensor. The recesses are plugged with clear resin corks as detailed on OSF.

8 These conditions were not contiguous; two other conditions were completed, one between VR2 baseline and extinction and one between extinction and VR2 return to baseline but are not described nor data presented here as presentation is planned for another manuscript. We also had an apparatus error early in the initial baseline. Therefore we do not present data during acquisition or transitions. Rats continued self-administration beyond the data reported here.

9 Other details are available in supplemental files on OSF.

10 With inferential statistical analysis of small samples, we run the risk of a Type II error rather than Type I error (Sullivan et al., 2016). In other words, with small sample sizes, we risk failing to find an effect when there may be one, rather than finding an effect where there is none.

11 Transforming the data into rank orders to reflect the non-normal distributions did not change the outcome of the results. A parametric ANOVA on the unranked data produced the same results; these results can be found in Prism files on OSF.

12 We give a subset of these tests in this report. All follow up tests can be found on OSF.

13 Shelton and Nicholson (2022) describe a nose-only chamber for aerosolized *opioid* self-administration in rats. However, this chamber was not yet verified to sequester the aerosol in the port, opioid levels were not assessed, and rats received food training in between each opioid-self administration condition.

## Notes

https://osf.io/x2pqf/?view_only=775b55435b8e428f98e6da384ef7889d

### Competing Interest Statement

The authors have declared no competing interest.

### Summary of Updates

Minor wording changes, reference additions, minor change of text on figure

